# Medulla glutamatergic neurons control wake-sleep transitions

**DOI:** 10.1101/2021.03.07.434263

**Authors:** Sasa Teng, Fenghua Zhen, Jose Canovas Schalchli, Xinyue Chen, Hao Jin, Li Wang, Yueqing Peng

## Abstract

Sleep is a ubiquitous behavior in animal species. Yet, brain circuits controlling sleep remain poorly understood. Previous studies have identified several brain structures that promote sleep, but whether these structures are involved in sleep initiation or sleep maintenance remains largely unknown. Here we identified a population of glutamatergic neurons in the medulla that project to the preoptic area (POA), a prominent sleep-promoting region. Chemogenetic silencing of POA-projecting medulla neurons disrupts the transitions from wakefulness to Non-Rapid Eye Movement (NREM) sleep, whereas chemogenetic activation of these neurons promotes NREM sleep. Moreover, we show that optogenetic activation of medulla glutamatergic neurons or their projections in the POA reliably initiates long-lasting NREM sleep in awake mice. Together, our findings uncover a novel excitatory brainstem-hypothalamic circuit that controls the wake-sleep transitions.

## INTRODUCTION

Sleep and wakefulness are actively controlled by dedicated neural circuits in the brain (Scammell et al., 2017; Weber and Dan, 2016). Centered on the mutual inhibition between sleep-promoting and wakefulness-promoting circuits, a flip-flop model has been proposed to explain the mechanisms underlying sleep-wake switching (Saper et al., 2001; Saper et al., 2010). The discovery of the ascending reticular activating system (Moruzzi and Magoun, 1949), the orexin neurons (de Lecea et al., 1998; Sakurai et al., 1998), and further studies of neuromodulatory systems have greatly advanced our understanding of the neural circuits supporting wakefulness (Brown et al., 2012; Lee and Dan, 2012; Scammell et al., 2017). In contrast, the neural circuits controlling sleep processes are largely elusive. Several sleep-promoting regions have been identified, including the preoptic area (POA, particularly the ventrolateral preoptic area and median preoptic nucleus, or VLPO and MPO) (Alam et al., 2014; John and Kumar, 1998; Kroeger et al., 2018; Lu et al., 2000; Sherin et al., 1996), the parafacial zone (PZ) (Anaclet et al., 2014; Anaclet et al., 2012), and the basal forebrain (BF) (Xu et al., 2015). GABAergic neurons in these brain regions are sleep-active, and activation of these neurons promotes NREM sleep(Scammell et al., 2017). For instance, Chung et. al. demonstrated that TMN-projecting GABAergic neurons in the POA were both sleep active and sleep promoting (Chung et al., 2017). Sleep-active neurons are also found in a subset of GABAergic cortical interneurons that produce neuronal nitric oxide synthase (nNOS) (Gerashchenko et al., 2008; Morairty et al., 2013). In addition to the role of GABAergic neurons in sleep control, a recent study identified an excitatory circuit in the perioculomotor (PIII) region of the midbrain that promotes NREM sleep (Zhang et al., 2019). They found that CALCA-expressing glutamatergic PIII neurons project to the POA and the medulla, and optogenetic activation of these projections can promote NREM sleep.

The POA is the most intensively studied sleep-active and sleep-promoting region. Immunohistochemistry studies in rats showed the existence of sleep-active GABAergic neurons in the VLPO and a correlation between the number of c-Fos positive cells and the amount of NREM sleep (Lu et al., 2002; Sherin et al., 1996). Consistently, electrophysiological recordings demonstrated that the firing rate of VLPO sleep-active neurons correlates with the depth of NREM sleep (indicated by EEG delta power) (Alam et al., 2014; Szymusiak et al., 1998). Notably, VLPO neurons displayed lower activity in the wake-sleep transition period than in the subsequent sleep episode (discharge rates further increased significantly from light to deep NREM sleep) (Szymusiak et al., 1998). These findings suggest that VLPO neurons might be involved in sleep maintenance whereas other neurons may be responsible for sleep initiation.

Early transection studies suggest the existence of sleep-promoting neurons in the medullary brainstem (Villablanca, 2004). Indeed, recent studies identified a population of GABAergic neurons in the parafacial zone (PZ) as a medullary NREM sleep promoting center (Anaclet et al., 2014; Anaclet et al., 2012). In this study, we sought to identify novel medullary neurons that might contribute to the regulation of sleep behavior. We reasoned that neurons projecting to the POA could be involved in sleep initiation. In particular, we hypothesized that presynaptic excitatory neurons activate postsynaptic POA GABAergic neurons to initiate sleep states.

## RESULTS

### Medulla glutamatergic neurons project to the preoptic area

To help identify sleep-promoting neurons in the brainstem, we performed retrograde tracing by injecting retrograde virus carrying Cre-recombinase (AAVrg-Cre) in the POA in a reporter transgenic line (Ai9, Figure 1A). Four weeks after viral injection, we perfused and examined the whole brains by CUBIC clearing and rapid 3D imaging with light-sheet fluorescent microscopy (Susaki et al., 2015). The whole-brain retrograde tracing uncovered multiple brain regions that project to the POA, with the majority in the forebrain (Figure 1B). Among tdTomato-labeled cells, we identified a population of neurons in the ventrolateral medulla (VLM) (Figure 1B-C). The labeled cells were largely clustered in areas overlapping C1 adrenergic and A1 noradrenergic fields, lateral and dorsal to the lateral paragigantocellular nucleus (LPGi) (Chou et al., 2002). A few cells were distributed inside the LPGi. To examine the identity of POA-projecting VLM neurons, we used retrograde AAVrg-FLEX-syn-H2B-Ruby3-H2B-Clover3 to express two nucleus-localized fluorescent proteins: Clover3 in all retrograde transduced cells and Ruby3 in Cre-expressing subset cells (Figure 1D). To test our hypothesis of excitatory connections, we performed retrograde tracing by injecting AAVrg-FLEX-syn-H2B-Ruby3-H2B-Clover3 into the POA of Vglut2-Cre mice. Indeed, the majority (∼60%) of Clover-labeled cells were co-labeled by Ruby in the medulla (Figure 1E-F). In contrast, we observed Clover-labeled cells but not Ruby-labeled cells in the medulla when we repeated retrograde tracing experiments in GAD2-Cre mice (Figure 1E-F). Together, these results indicated that medullary glutamatergic neurons, but not GABAergic neurons project to the POA.

**Figure 1.**
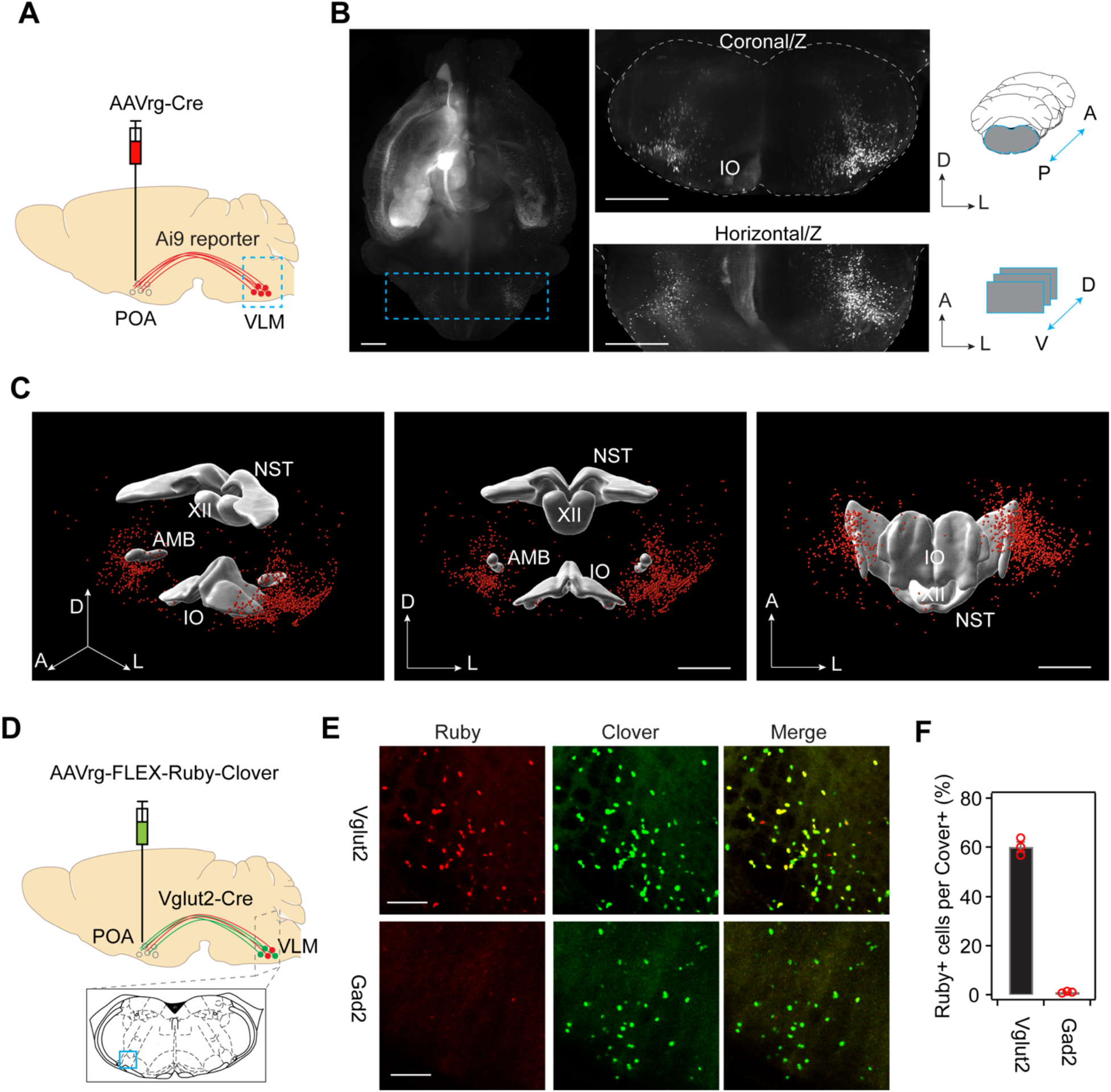
Medulla glutamatergic neurons project to the preoptic area. **A**, Schematic illustrating retrograde labelling experiments in Ai9 reporter mice injected with AAVrg-hSyn.Cre.WPRE.hGH in the preoptic area (POA). VLM, ventrolateral medulla. **B**, Left, maximum-intensity z stack of POA-projecting neurons in the whole mouse brain, cleared with CUBIC and imaged with light-sheet fluorescent microscopy. The dashed-line box illustrating the anterior and posterior boundaries used for reconstruction of coronal and horizontal views on the right. Right top, Maximum-intensity z stack of POA-projecting neurons in the optically sliced coronal sections of the brainstem. Right bottom, Maximum-intensity z stack of POA-projecting neurons in the optically sliced horizontal sections of the brainstem. Scale bars, 1 mm. **C**, Three-dimensional reconstruction of POA-projecting medulla neurons in the perspective view, the front view, and the ventral view, respectively. A, anterior; D, dorsal; L, lateral. NST, nucleus of the solitary tract; XII, hypoglossal nucleus; AMB, nucleus ambiguus; IO, inferior olivary complex. Scale bars, 1 mm. **D**, Schematic of retrograde tracing in Vglut2-Cre mice injected with AAVrg-H2B-Clover3-FLEX-H2B-Ruby3 in the POA. Bottom, coronal section (Bregma -6.84 mm) showing the VLM. **E**, Representative image of Cre-expressing cells (Ruby) and all POA-projecting cells (Clover) in the VLM (blue box in schematic) in Vglut2-Cre (upper) and Gad2-Cre (lower) animals. Scale bars, 100 μm. **F**, Quantitation of percentage of Cre-expressing cells among all POA-projecting cell (n = 3 animals for Vglut2-Cre, n = 3 animals for Gad2-Cre).

The POA contains different cell types of neurons. For instance, about 85% of the neurons in the VLPO contain the inhibitory neurotransmitters galanin and GABA (Sherin et al., 1998; Sherin et al., 1996), which are sleep-active and sleep-promoting (Alam et al., 2014; John and Kumar, 1998; Kroeger et al., 2018; Lu et al., 2000; Sherin et al., 1996). To examine whether medulla neurons directly synapse with POA GABAergic neurons, we used a Cre-dependent monosynaptic retrograde viral reporter to study their connections (Callaway and Luo, 2015; Reardon et al., 2016). In GAD2-Cre mice, we injected AAV-FLEX-G(N2C)-mKate (Reardon et al., 2016) and AAV-FLEX-TVA-mCherry into the POA. Two weeks after AAV injection, we injected RABV-∆G -GFP-EnvA (Reardon et al., 2016) into the same POA region. Seven to ten days after rabies injection, mice were perfused to examine GFP expression in the brain. As expected, we identified GFP labeled neurons in the VLM (Figure 2), which indicated that VLM neurons are directly synapse with GABAergic neurons in the POA.

**Figure 2.**
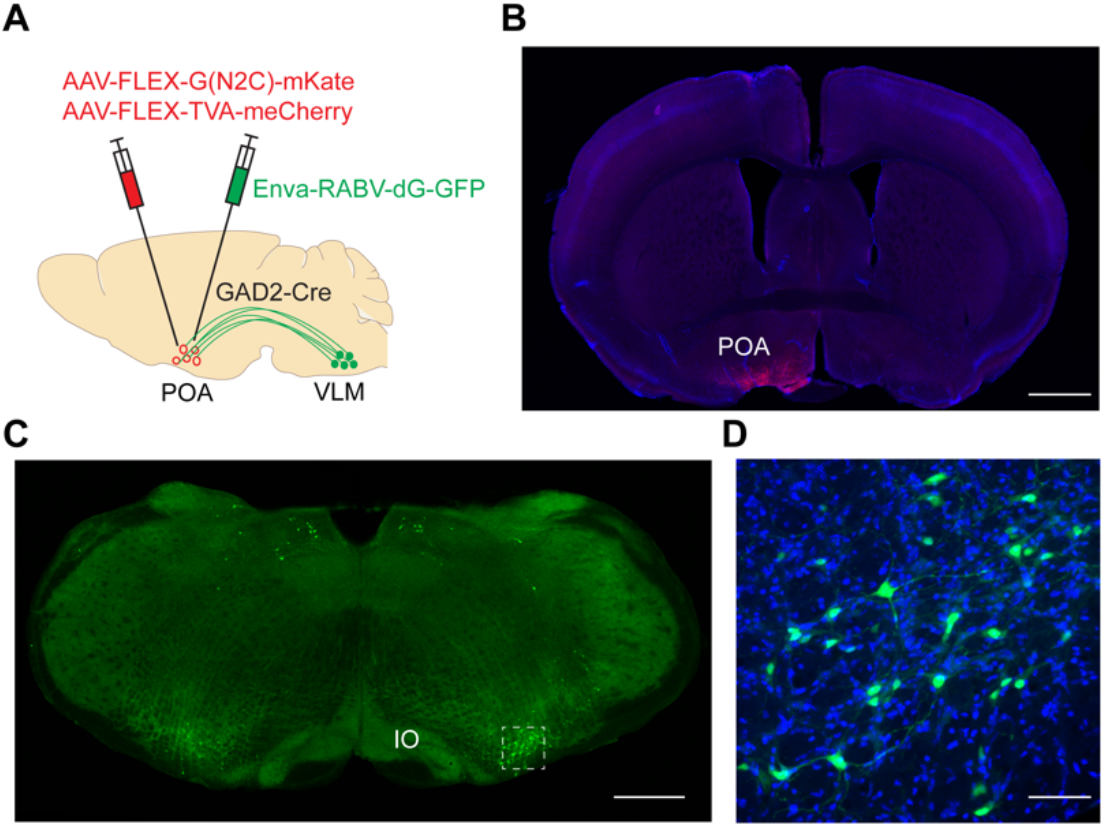
Monosynaptic retrograde tracing from the POA to VLM. **A**, Schematic of experimental design. GAD2-Cre animals were infected with AAV-FLEX-G(N2C)-mKate and AAV-FLEX-TVA-mCherry, followed by infection with Enva-RABC-∆G-GFP in the POA. **B**, Starter cells in the POA labeled by mKate/mCherry (red). Blue, DAPI. Scale bar, 1 mm. **C**, Rabies-labeled cells in the VLM indicated by GFP expression (green). Scale bar, 0.5 mm. **D**, Enlarged view. Green, GFP; Blue, DAPI. Scale bar, 50 μm.

### POA-projecting medulla neurons are required for wake-sleep transitions

Next, we conducted both loss-of-function and gain-of-function experiments to determine the role of the VLM-POA circuitry in sleep behavior. To test the necessity, we chemogenetically silenced POA-projecting VLM neurons and then examine its effects on wake-sleep transitions. To selectively target POA-projecting glutamatergic neurons in the VLM, we bilaterally injected retrograde AAV expressing Cre recombinase (AAVrg-Cre) into the POA, and bilaterally injected AAV carrying Cre-dependent inhibitory DREADDs (AAV9-DIO-hM4D(Gi)) into the VLM (Figure 3A). As shown in Figure 1, retrograde AAV injected to the POA predominately labels glutamatergic neurons in the VLM. After two weeks of recovery, we treated animals with clozapine-N-oxide (CNO, 1 mg/kg) or saline and examined their effects on sleep behavior (Figure 3B). We reasoned that if POA-projecting VLM neurons are required for the transitions from wakefulness to sleep, we expect to observe less NREM sleep and more wake time after CNO treatment. Since NREM sleep is often followed by REM sleep, we also reasoned that less NREM sleep could lead to less REM sleep. Indeed, silencing VLM neurons significantly promoted wakefulness and suppress both NREM sleep and REM sleep (Figure 3C). Further analysis demonstrated that decreased NREM and REM sleep were caused by decreased bout numbers of NREM and REM sleep, while the bout durations of NREM and REM sleep remain unchanged (Figure 3D-E). In control experiments, CNO treatment (1 mg/kg) has no significant effect on sleep behavior in the wild type mice without DREADDs expression (data not shown). Together, these results indicate that POA-projecting VLM neurons are required for the transitions from wakefulness to NREM sleep.

**Figure 3.**
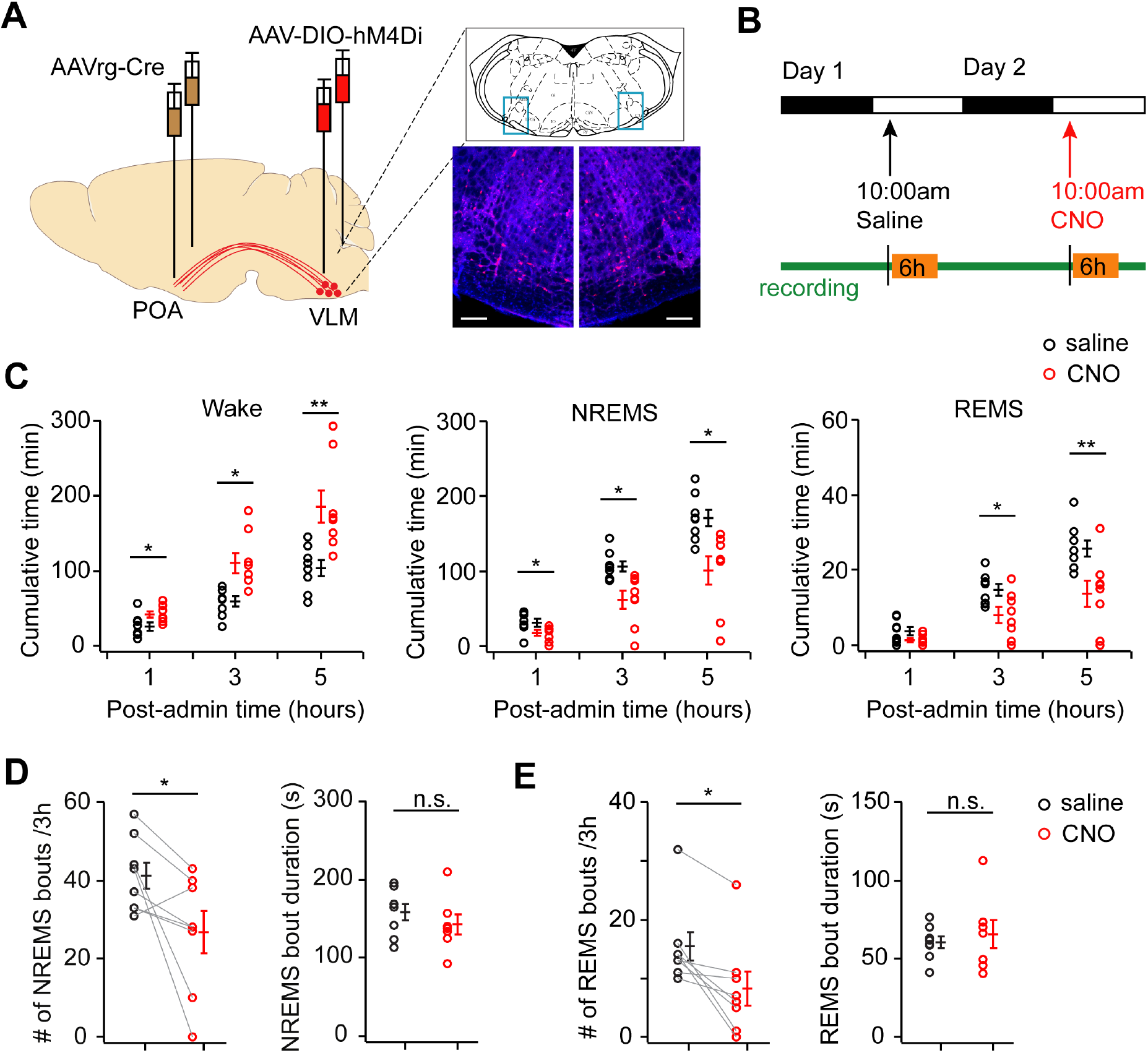
Silencing medulla glutamatergic neurons disrupts the transitions from wakefulness to sleep. **A**, Schematic of chemogenetic silencing of POA-projecting medulla neurons. AAVrg-Cre was bilaterally injected in the POA and AAV9-hSyn-DIO-hM4D(Gi)-mCherry was bilaterally injected in the VLM. Right, Fluorescence images of coronal sections in the VLM (blue boxes above) showing bilateral expression of inhibitory DREADDs. Red, mCherry, Blue, DAPI. Scale bars, 100 μm. **B**, Schematic of drug administration and sleep recording. **C**, Quantitation of the total wake, NREM sleep, and REM sleep time following CNO (1 mg/kg, i.p. red circles) and saline (black circles) treatment (n = 8 mice, paired t-test). **D**, Quantitation of bout numbers and bout durations of NREM sleep within 3 hours after CNO and saline treatment (n = 8 animals, paired t-test). **E**, Quantitation of bout numbers and bout durations of REM sleep within 3 hours after CNO and saline treatment (n = 8 animals, paired t-test, n.s. no significance, * P<0.05, ** p<0.01).

### POA-projecting medulla neurons are sufficient to promote NREM sleep

We then examined the sufficiency of POA-projecting VLM neurons in wake-sleep transitions by conducting gain-of-function experiments. Using a similar dual viral injection strategy, we chemogenetically activated POA-projecting VLM neurons with and examined its effect on sleep behavior. We bilaterally injected AAVrg-Cre into the POA, and unilaterally injected AAV carrying Cre-dependent excitatory DREADDs (AAV8-DIO-hM3D(Gq)) into the VLM (Figure 4A). After two weeks of recovery, we treated animals with CNO (1 mg/kg) and saline while recording sleep behavior (Figure 4B). We found that chemogenetic activation of POA-projecting VLM neurons significantly increased the amount of time animals spent in NREM sleep, accompanied with the decrease of wake time, (Figure 4). We observed no significant change of the total REM sleep time after CNO treatment (Figure 4C). Further analysis showed that the increase of NREM sleep time was largely due to the increase of NREM sleep bouts, but not the bout durations of NREM sleep (Figure 4D). These results indicate that chemogenetic activation of POA-projecting neurons in the VLM is sufficient to promote NREM sleep.

**Figure 4.**
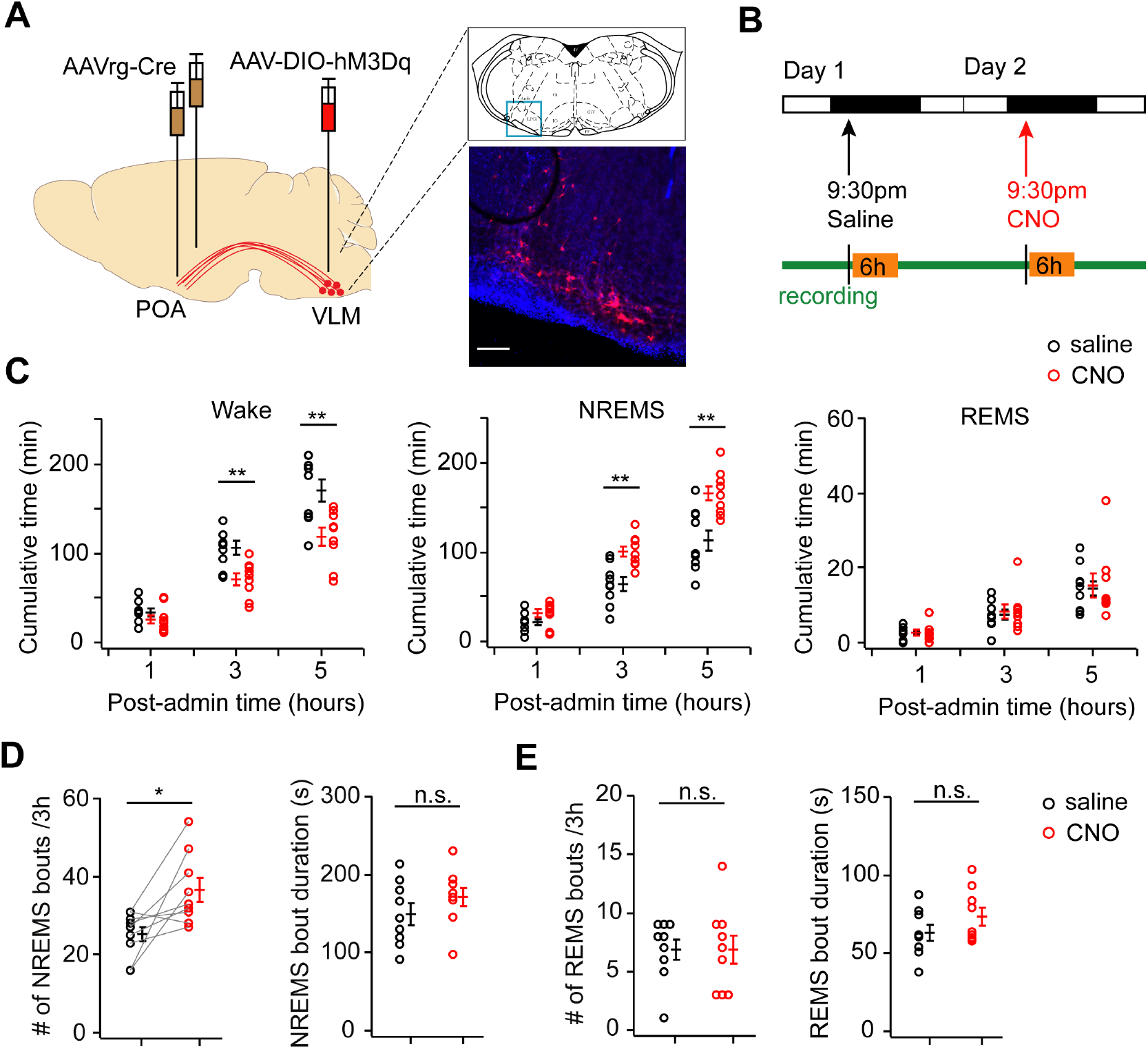
Chemogenetic activation of medulla glutamatergic neurons promotes NREM sleep. **A**, Schematic of chemogenetic activation of POA-projecting medulla neurons. AAVrg-Cre was bilaterally injected in the POA, AAV8-hSyn-DIO-hM3D(Gq)-mCherry was unilaterally injected in the VLM. Right, Fluorescence image of a coronal section in the VLM (blue box above) showing expression of excitatory DREADDs. Red, mCherry, Blue, DAPI. Scale bar 100 μm. **B**, Schematic of drug administration and sleep recording. **C**, Quantitation of the total wake, NREM sleep, and REM sleep time after CNO (1 mg/kg, i.p.) and saline treatment (n = 9 animals, paired t-test). **D**, Quantitation of bout numbers and bout durations of NREM sleep within 3 hours after CNO and saline treatment (n = 9 animals, paired t-test). **E**, Quantitation of bout numbers and bout durations of REM sleep within 3 hours after CNO and saline treatment (n=9 animals, paired t-test, n.s. no significance, * P<0.05, ** p<0.01).

### Medulla glutamatergic neurons control wake-sleep transitions

The transitions from wakefulness to NREM sleep occur in the time scale of seconds. Due to the slow process of drug administration, chemogenetic manipulation lacks temporal resolutions to track each wake-sleep transition. To address this problem, we then used optogenetics to activate the VLM neurons and examine behavioral output. To directly target VLM glutamatergic neurons, we stereotaxically injected AAV1-DIO-ChR2-eYFP into the VLM in Vglut2-Cre mice and implanted an optic fiber above the injection site (Figure 5A). Upon the offset of light stimulation (20 Hz, 2 min), we observed rapid transitions from wakefulness to NREM sleep (Figure 5B).

**Figure 5.**
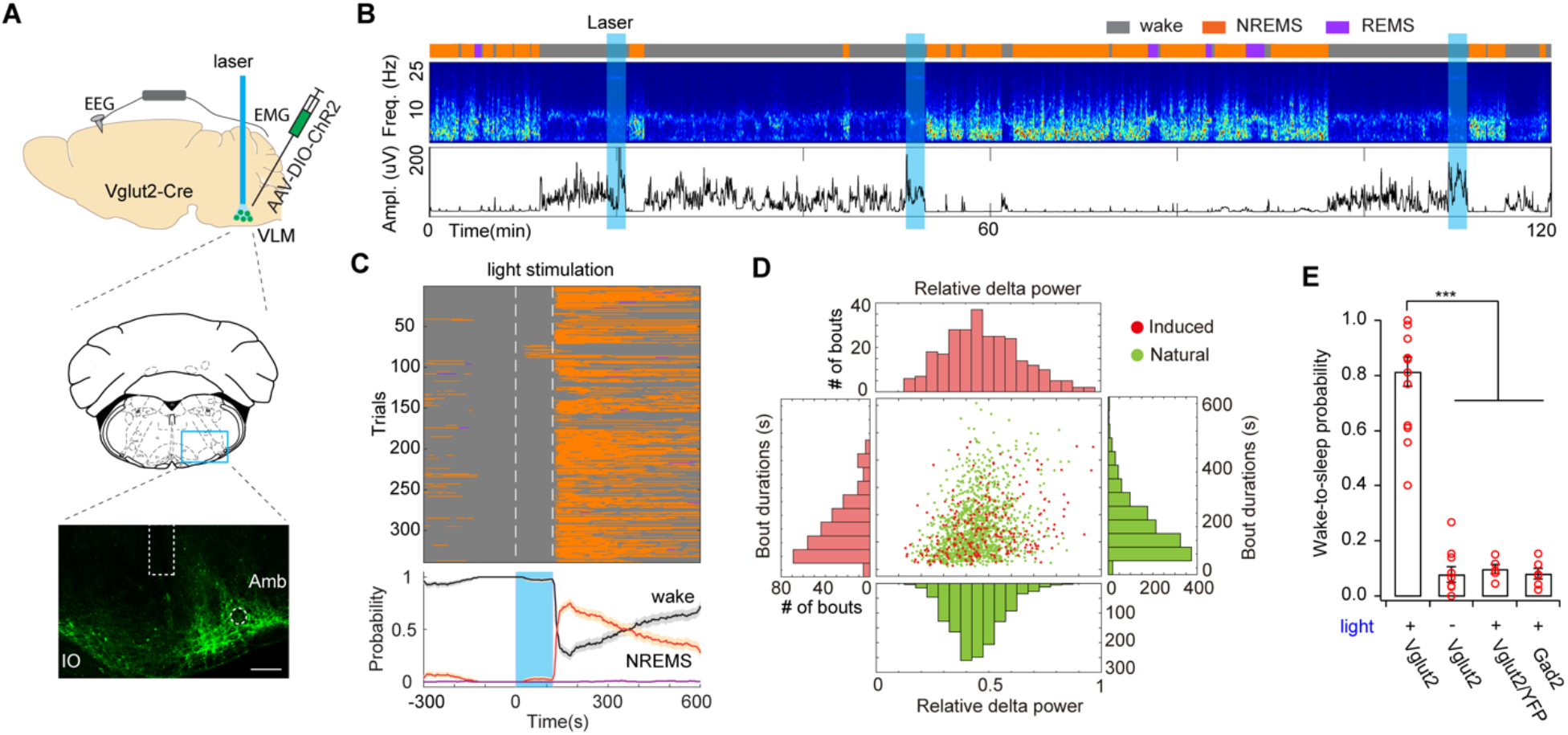
Optogenetic activation of medulla glutamatergic neurons initiates NREM sleep in awake animals. **A**, Schematic of optogenetic experiment. Middle, mouse brain coronal section (Bregma -6.84 mm). Bottom, fluorescence image of VLM (blue box above) in a Vglut2-Cre mouse injected with AAV expressing ChR2-eYFP (green). Dashed box indicates the placement of an optic fiber. Amb, nucleus ambiguus; IO, inferior olivary complex. Scale bar 100 μm. **B**, Representative EEG spectrogram and EMG trace in a 2-h session showing the induction of NREM sleep after photostimulation (20 Hz, 2 min, blue shade) of medulla glutamatergic neurons. Brain states: orange, NREM sleep; purple, REM sleep; gray, wake. **C**, Brain states before, during and after laser stimulation (white dash lines, 20 Hz, 2 min) in awake animals (n = 14). Bottom, probability of wake (black), NREM sleep (orange), and REM sleep (purple). Shading, 95% bootstrap confidence intervals. **D**, Scatter plots and distributions of bout durations and relative delta power in optogenetic-induced and natural NREM sleep (n = 14 animals). **E**, Quantification of transition probability from wakefulness to NREM sleep induced by optogenetic activation (n = 14 for Vglut2-Cre with light, n = 10 for Vglut2-Cre without light, n = 5 for Vglut2-Cre/YFP, n = 6 for Gad2-Cre, Mann-Whitney *U*-test, *** P<0.001).

To quantitatively examine the probability of optogenetically induced transitions from wakefulness to sleep, we applied video-based closed-loop optogenetic stimulation in awake animals. An IR-camera was used to real-time track animal’s movement and laser stimulation was automatically triggered by wakefulness. We found that optogenetic activation of VLM glutamatergic neurons reliably initiated NREM sleep in awake mice with a transition probability of ∼80% (Figure 5C, E). Notably, optogenetic-induced NREM sleep episodes lasted a few minutes, with a distribution of bout durations similar to that in natural sleep (Figure 5D). Moreover, sleep depth indicated by the delta power showed no significant difference between optogenetic-induced and natural NREM sleep episodes (Figure 5D). To our knowledge, this is the first study that optogenetic stimulation can initiate natural-like NREM sleep. In Vglut2-Cre mice, we performed two control experiments: 1) no light stimulation in ChR2-expressing mice, 2) light stimulation in YFP-expressing mice (AAV1-DIO-eYFP). As expected, we observed low spontaneous but not significant wake-sleep transitions following photostimulation (Figure 5E). As another control, we performed similar optogenetic experiments in GAD2-Cre mice. No significant transitions from wakefulness to NREM sleep were observed upon activation of VLM GABAergic neurons in awake animals (Figure 5E). Together, these results indicate that optogenetic activation of VLM glutamatergic neurons can reliably initiate the transition from wakefulness to NREM sleep.

A notable phenomenon in this study was that NREM sleep occurred following the offset of laser stimulation rather than the onset. There are two possibilities: 1) it takes a few minutes of light stimulation to induce wake-sleep transitions, and the timing effect is a coincidence due to the stimulation protocol we used; 2) nonspecific activation prevents animals falling asleep during the stimulation period. To test the first possibility, we activated the VLM glutamatergic neurons with different durations of laser stimulation (30 s and 1 min). We found that wake-sleep transitions similarly occurred following the offset of laser stimulation (Figure 6A-B). Furthermore, shorter durations of stimulation triggered less wake-sleep transitions indicated by slightly lower probabilities (Figure 6C). These results excluded the first possibility and suggested that nonspecific activation might be the reason, either caused by viral spread to the adjunct regions (see below) or the heterogeneity of glutamatergic neurons in the VLM.

**Figure 6.**
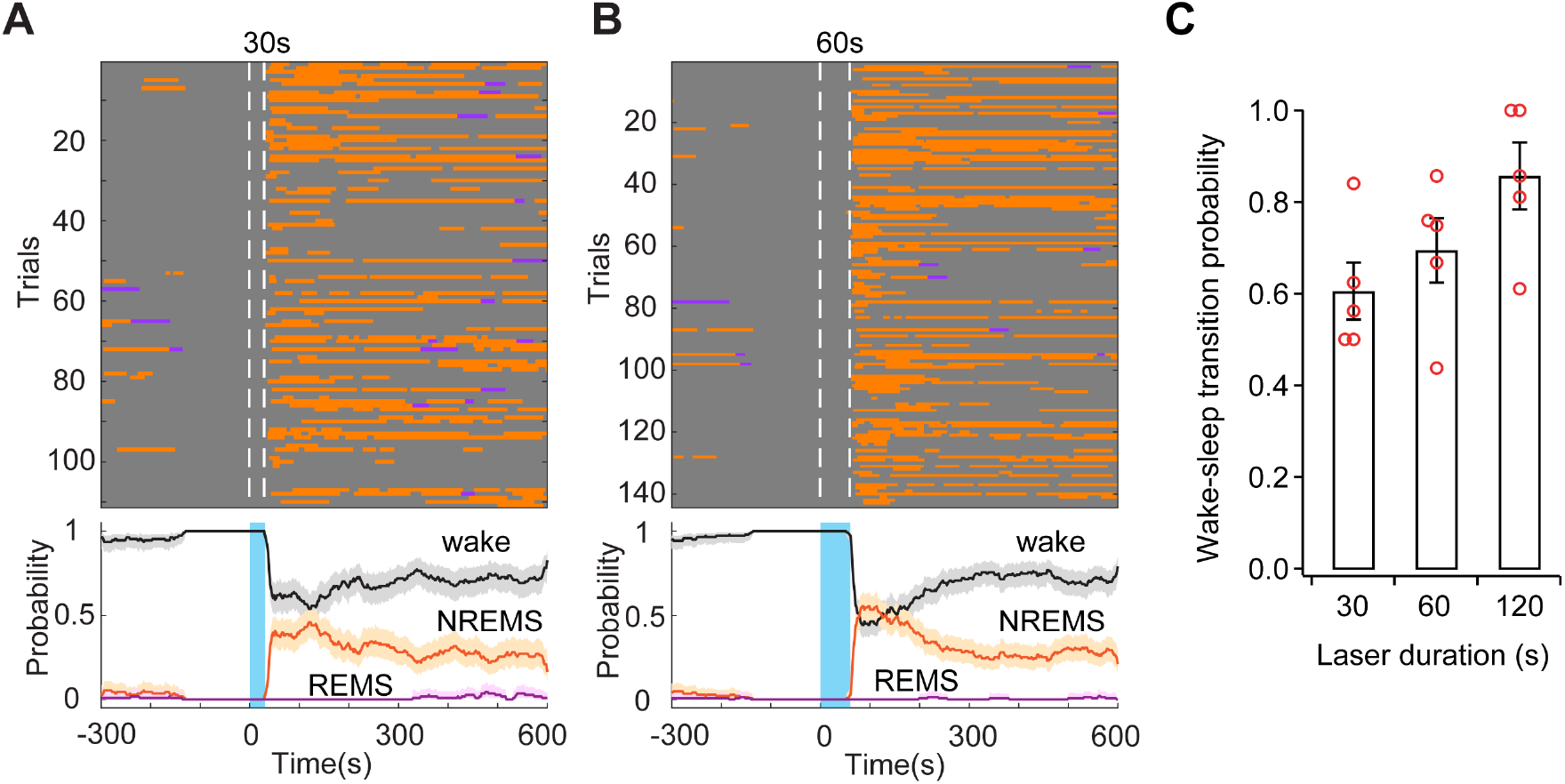
Optogenetic stimulation with different durations. **A**, Brain states (upper) and probability (bottom) of wake (gray), NREM sleep (orange), and REM sleep (purple) in trials with wakefulness before laser stimulation (30 s, 20 Hz) from 5 animals. Shading, 95% bootstrap confidence intervals. **B**, Brain states (upper) and probability (bottom) of wake, NREM sleep, and REM sleep in trials with wakefulness before laser stimulation (60 s, 20 Hz). **C**, Quantitation of wake-sleep transition probability with different durations of stimulation (n = 5 Vglut2-Cre mice).

Next, we examined the effect of optogenetic stimulation in sleeping mice by using a fixed-interval (45 min) light stimulation. In trials with sleep (both NREM sleep and REM sleep) before laser stimulation, we found that activation of VLM glutamatergic neurons strongly interrupted animal’s sleep and triggered immediate locomotor activity (Figure 7). Interestingly, in the majority of these trials, animals were able to switch back to NREM sleep when the light was off. We also noticed optogenetic-induced locomotor activity in some awake animals. We speculated that the locomotion behavior might be caused by viral spread and non-specific stimulation in a motor area in the lateral paragigantocellular nucleus (LPGi), which is adjacent to the target brain region. Capelli et. al. showed that optogenetic activation of glutamatergic neurons in the LPGi elicits full body locomotion (Capelli et al., 2017). This optogenetic-induced nonspecific locomotion might awaken animals from sleep. In addition, this locomotion effect might also cause disturbances and prevent animals from falling asleep during the photostimulation period, which could be the reason why optogenetic-induced wake-sleep transitions happen following the offset of laser stimulation.

**Figure 7.**
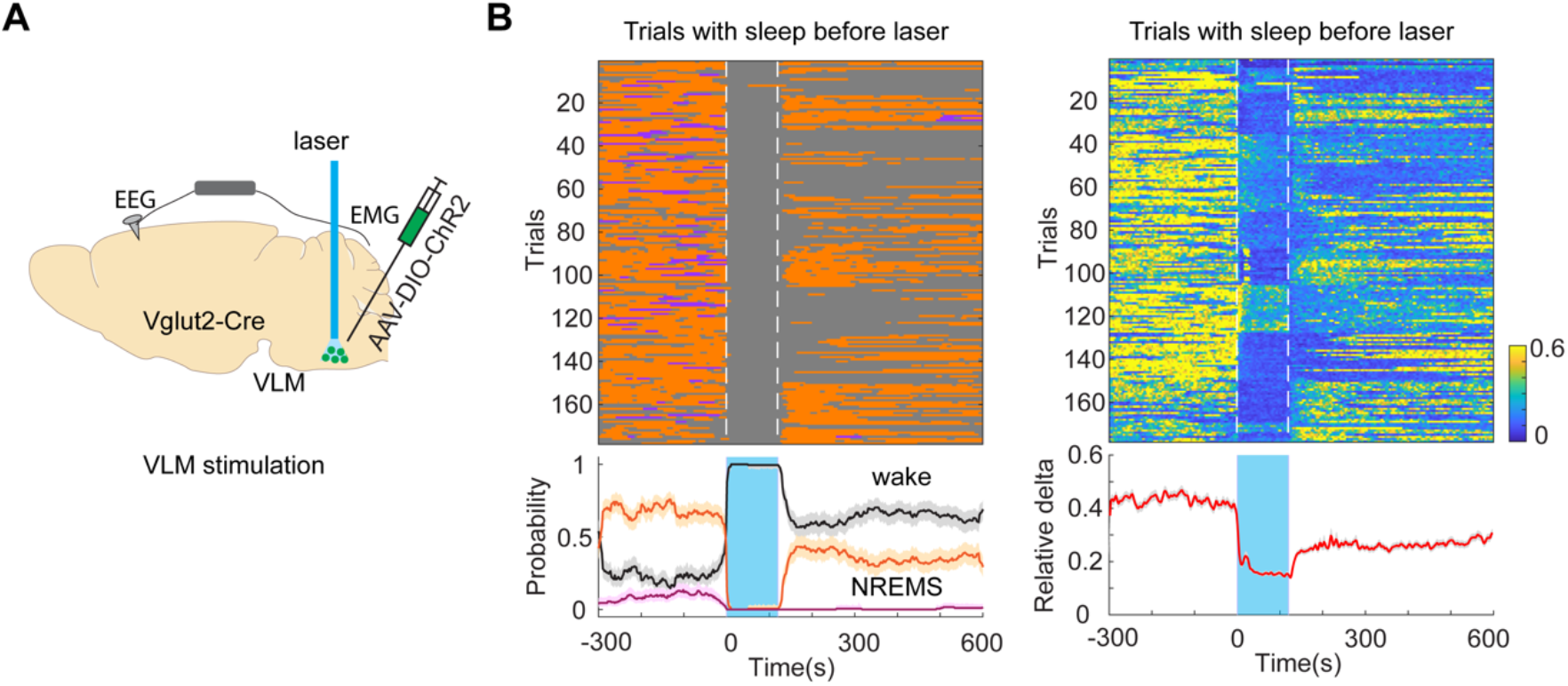
Optogenetic activation of VLM glutamatergic neurons in sleeping mice. **A**, schematic of experimental design. **B**, Optogenetic activation of VLM glutamatergic neurons interrupted sleep in sleeping animals (n = 14 animals). Left, brain states (upper) and probability (bottom) before, during, and after light stimulation (20 Hz, 2min), Shading of traces, 95% bootstrap confidence intervals. Right, relative delta power in each trial. Shading of trace, s.e.m.

To avoid nonspecific activation, and more importantly to specifically activate the VLM-POA pathway, we performed terminal stimulation in Vglut2-Cre animals. To identify the projection patterns of VLM glutamatergic neurons in the POA, we first performed anterograde tracing by unilaterally injecting AAV-FLEX-tdTomato into the VLM of Vglut2-Cre animals (Figure 8A). we found that TdTomato-labelled neurons projected to several regions in the POA with particularly dense innervation into the VLPO (Figure 8B). Then, we injected AAV1-DIO-ChR2-eYFP into the VLM and implanted an optic fiber above the VLPO (Figure 8C). We found that optogenetic activation of VLM glutamatergic terminals in the POA initiated NREM sleep in awake animals following the onset of laser stimulation (Figure 8D). Further analysis demonstrated that the bout durations were shorter in optogenetic-induced NREM sleep, compared to that in natural NREM sleep, whereas the delta power was not significantly different between groups (Figure 8E). Furthermore, the wake-sleep transition probability in terminal stimulation was lower than that in VLM soma stimulation (Figure 8D). In addition, we did not observe significant locomotor activity and sleep interruption upon terminal stimulation in sleeping mice (Data not shown). Together, these results indicated that optogenetic activation of the VLM-POA circuitry can initiate the transitions from wakefulness to NREM sleep.

**Figure 8.**
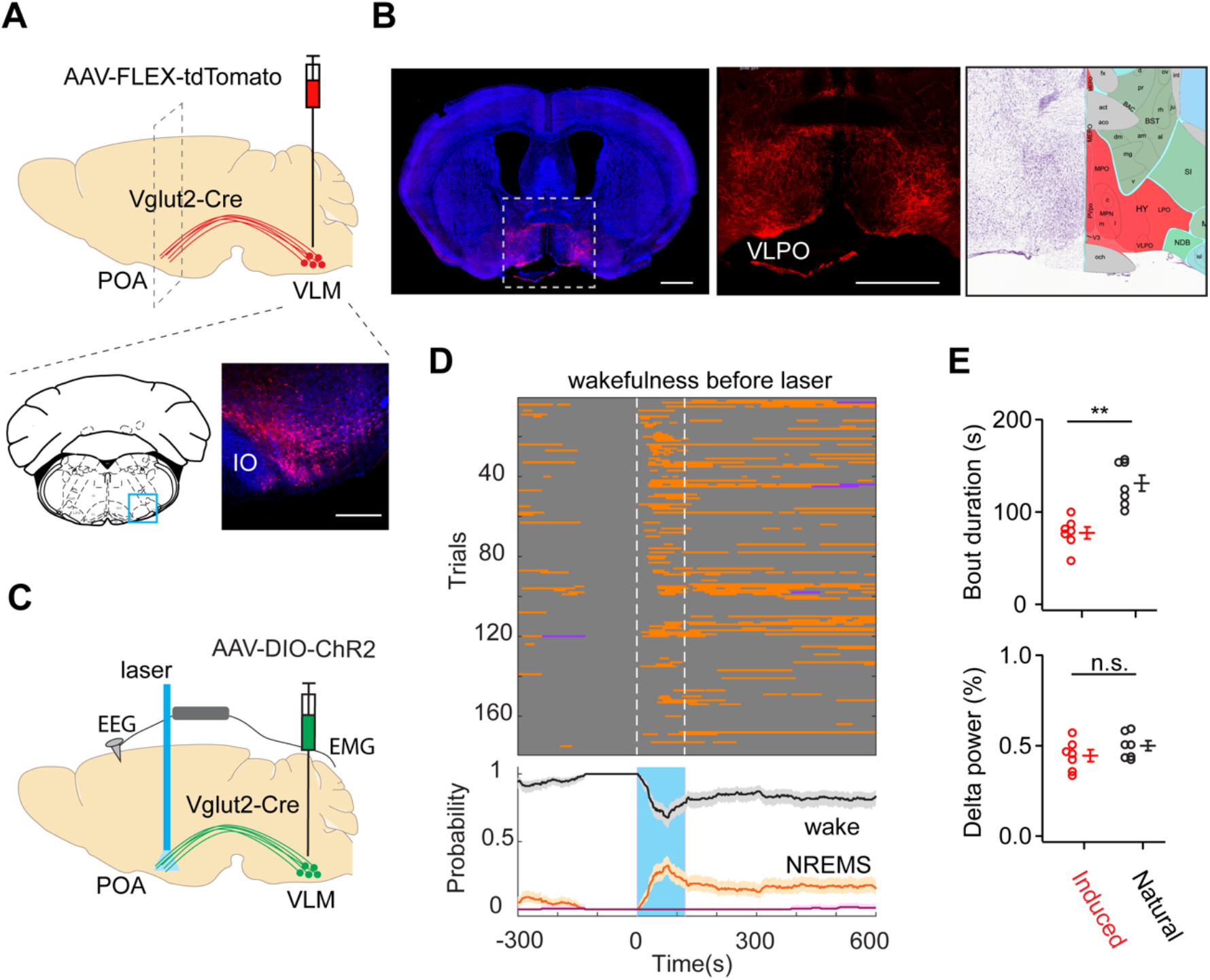
Optogenetic activation of VLM glutamatergic terminals in the POA promotes the transition from wakefulness to NREM sleep. **A**, Schematic of anterograde tracing experiment. AAV1-CAG-FLEX-tdTomato (red) was unilaterally injected in the VLM (bottom) of Vglut2-Cre mice. Blue DAPI. Scale bar, 0.2 mm. **B**, fluorescence image of a coronal section (Bregma 0) illustrating axonal terminals (red) in the POA (enlarged in the middle). Blue DAPI. Scale bar, 1 mm. Right, diagram of the POA from the Allen brain atlas. **C**, Schematic of optogenetic experiment. **D**, Brain states (upper) and probability (bottom) before, during, and after laser stimulation of VLM glutamatergic terminals in the POA of awake animals (n = 7). **E**, Bout durations (upper) and relative delta power (bottom) in optogenetic-induced and natural NREM sleep (n = 7 animals, ** p<0.01, paired t-test).

## DISCUSSION

In this study, we identified a population of glutamatergic neurons in the medulla that project to the POA, a prominent sleep-promoting center. Using chemogenetic manipulation, we demonstrated that these POA-projecting medulla neurons are both necessary and sufficient for the transitions from wakefulness to sleep. Furthermore, optogenetic activation of medulla glutamatergic neurons or their axonal terminals in the POA robustly initiates NREM sleep in awake animals. Together, our results uncovered a novel excitatory brainstem-hypothalamic circuit that control the transitions from wakefulness to NREM sleep (i.e., sleep initiation). Notably, our finding of sleep-promoting excitatory glutamatergic cells in the medulla differs from the sleep-promoting GABAergic neurons previously reported in the POA (Alam et al., 2014; John and Kumar, 1998; Lu et al., 2000; Sherin et al., 1996), BF (Xu et al., 2015), PZ (Anaclet et al., 2014; Anaclet et al., 2012), and nNOS+ cortical neurons (Gerashchenko et al., 2008; Morairty et al., 2013).

REM sleep-controlling GABAergic neurons have been identified in the ventral medulla(Weber et al., 2015), which is adjunct to the glutamatergic neurons in this study. In our optogenetic experiments, we did observe that activation of VLM GABAergic neurons in some sleeping animals promoted REM sleep (data not shown), possibly due to the viral spread into the adjunct region. Previous studies also show that glutamatergic neurons in the ventromedial medulla regulate REM sleep and motor atonia (Chen et al., 2017; Vetrivelan et al., 2009). In both chemogenetic and optogenetic experiments, we did not observe significant changes of REM sleep following the activation of VLM glutamatergic neurons. Our retrograde tracing data demonstrated the POA-projecting glutamatergic neurons are located more laterally (Figure 1), compared to the neurons involved in REM sleep atonia (i.e., in a restricted region within the ventromedial medulla, termed supraolivary medulla). Intriguingly, the co-existence of REM sleep-promoting and NREM sleep-promoting neurons in the medulla implies the possible interaction among these neurons and highlights the importance of the medulla in sleep regulation.

Interestingly, NREM sleep initiated by optogenetic activation of VLM neurons usually lasted several minutes and showed similar bout durations to that in natural sleep. This sleep phenomenon is novel and different from other optogenetic studies in the POA (Chung et al., 2017), and the PIII (Zhang et al., 2019), where the effect of optogenetic activation quickly disappears when light stimulation is off. It suggests that sustained stimulation is required for the POA or PIII neurons to promote and more importantly to maintain NREM sleep, whereas it is not necessary to maintain NREM sleep in our case. Our data suggest the VLM glutamatergic neurons may function as an initiator to trigger the wake-sleep transition process, and subsequently their activity may not be required for sleep maintenance. Compared to soma stimulation in the VLM, we noticed that terminal stimulation in the POA was less effective to initiate NREM sleep, considering both lower wake-sleep transition probability and shorter bout durations (Figure 8). These results imply that VLM neurons may project to other brain areas to orchestrate the sleep initiation process. Further studies on inputs and outputs of VLM neurons will shed light on mechanistic understanding of this process.

## ACKNOWLEDGEMENTS

We thank Charles Zuker at Columbia University for his support in the early stage of the study. We thank Xiaoke Chen at Stanford University for the gift of the retrograde virus; we acknowledge the Cellular Imaging Core at the Columbia Zukerman Institute for access to their facility. We also thank Xiaoke Chen and Charles Zuker for helpful discussions. This work was supported by the startup fund from Columbia University.

## AUTHOR CONTRIBUTIONS

S.T. and Y.P. designed the study, carried out the experiments, and analyzed data, F.Z. performed histology and circuit tracing experiments. J.C.S. performed behavioral experiments. X.C. performed data analysis. H.J. performed rabies-based tracing experiments. L.W. performed CUBIC experiments. Y.P. and S.T. wrote the paper.

## DECLARATION OF INTERESTS

Authors declare that they have no competing interests.

## METHODS

### Animals

All procedures were carried out in accordance with the US National Institute of Health (NIH) guidelines for the care and use of laboratory animals, and approved by the Animal Care and Use Committees of Columbia University. Both male and female adult mice which are older than 8 weeks of age were used for all experiments. The following mouse lines were used in the current study: C57BL/6J (JAX 000664), VGlut2-IRES-Cre (JAX 028863), Gad2-IRES-Cre (JAX 0101802), Ai9 (JAX 007909). Mice were housed in 12-hour light-dark cycles (lights on at 07:00 am and off at 07:00 pm) with free access to food and water.

### Viral constructs

AAV1-EF1α-double floxed-hChR2(H134R)-EYFP-WPRE-HGHpA, AAV1-Ef1α-DIO EYFP, AAV8-hSyn-DIO-hM3D(Gq)-mCherry, AAV9-hSyn-DIO-hM4D(Gi)-mCherry, AAVrg-hSyn-Cre-WPRE-hGH, AAV1-CAG-FLEX-tdTomato were obtained from Addgene. AAV1-EF1a-FLEX-TVA-mCherry from UNC vector core, AAV1-FLEX-2A-G(N2C)-mKate and RABV-∆G - GFP-EnvA, a gift from Charles Zuker. AAVrg-H2B-Clover3-FLEX-H2B-Ruby3, a gift from Xiaoke Chen at Stanford University.

### Stereotaxic surgery

Mice were anaesthetized with a mixture of ketamine and Xylazine (100 mg kg^-1^ and 10 mg kg^-1^, intraperitoneally), then placed on a stereotaxic frame with a closed-loop heating system to maintain body temperature. After asepsis, the skin was incised to expose the skull and a small craniotomy (∼0.5 mm in diameter) was made on the skull above the regions of interest. A solution containing 100-200 nl viral construct was loaded into a pulled glass capillary and injected into the target region using a Nanoinjector (WPI). Optic fibers (0.2 mm diameter, 0.39 NA, Thorlabs) were implanted into the target region with the tip 0.4 mm above the virus injection site for optogenetic manipulation. For EEG and EMG recordings, a reference screw was inserted into the skull on top of the cerebellum. EEG recordings were made from two screws on top of the cortex 1 mm from midline, 1.5 mm anterior to the bregma and 1.5 mm posterior to the bregma, respectively. Two EMG electrodes were bilaterally inserted into the neck musculature. EEG screws and EMG electrodes were connected to a PCB board which was soldered with a 5-position pin connector. All the implants were secured onto the skull with dental cement (Lang Dental Manufacturing). After surgery, the animals were returned to home-cage to recover for at least two weeks before any experiment.

For retrograde tracing, 150-200 nl AAVrg-hSyn.Cre.WPRE.hGH was unilaterally or bilaterally injected into the ventrolateral preoptic area (VLPO, Bregma 0.1 mm, lateral 0.9 mm, ventral 5.4 mm) of Ai9 mice, or 150-200 nl AAVrg-H2B-Clover3-FLEX-H2B-Ruby3 was unilaterally or bilaterally injected into the VLPO of Vglut2-Cre or Gad2-Cre mice. For rabies tracing, 200 nl mix of AAV-FLEX-G(N2C)-mKate and AAV-FLEX-TVA-mCherry (1:1) was unilaterally injected into the VLPO of Gad2-Cre mice. Two weeks after AAV injection, 200 nl Enva-RABC-∆G-GFP was unilaterally injected into the same VLPO. For anterograde tracing, 50 nl AAV1-CAG-FLEX-tdTomato was unilaterally injected into the ventrolateral medulla (VLM, Bregma -6.9 mm, lateral 1.1 mm, ventral 5.6 mm) of Vglut2-Cre mice. For optogenetic activation experiments, 200 nl AAV1-EF1α-double floxed-hChR2(H134R)-EYFP-WPRE-HGHpA was unilaterally injected into the VLM with optic fiber 0.4 mm on top of the viral injection site in Vglut2-cre mice. For terminal stimulation, 200 nl AAV1-EF1α-double floxed-hChR2(H134R)-EYFP-WPRE-HGHpA was unilaterally injected into the VLM of Vglut2-Cre mice, and an optic fiber was implanted in the POA. For chemogenetic inhibition, 200 nl AAV9-hSyn-DIO-hM4D(Gi)-mCherry was bilaterally injected in VLM, 200 nl AAVrg-hSyn.Cre.WPRE.hGH was bilaterally injected into the VLPO of C57BL/6J mice. For chemogenetic activation, 200 nl AAV8-hSyn-DIO-hM3D(Gq)-mCherry was unilaterally injected into the VLM, 200 nl AAVrg-hSyn.Cre.WPRE.hGH was bilaterally injected into the VLPO of C57BL/6J mice. The ventral coordinates listed above are relative to the pial surface.

### Sleep recording

Mouse sleep behavior was monitored using EEG and EMG recording along with an infrared video camera at 30 frames per second. Recordings were performed for 24-48 hours (light on at 7:00 am and off at 7:00 pm) in a behavioral chamber inside a sound attenuating cubicle (Med Associated Inc.). Animals were habituated in the chamber for at least 4 hours before recording. EEG and EMG signals were recorded, bandpass filtered at 0.5-500 Hz, and digitized at 1017 Hz with 32-channel amplifiers (TDT, PZ5 and RZ5D or Neuralynx Digital Lynx 4S). Spectral analysis was carried out using fast Fourier transform (FFT) over a 5 s sliding window, sequentially shifted by 2 s increments (bins). Brain states were semi-automatically classified into wake, NREM sleep, and REM sleep states using a custom-written Matlab program (wake: desynchronized EEG and high EMG activity; NREM: synchronized EEG with high-amplitude, delta frequency (0.5–4 Hz) activity and low EMG activity; REM: high power at theta frequencies (6–9 Hz) and low EMG activity). Semi-auto classification was validated manually by trained experimenters. Relative delta power was calculated by dividing the delta power in the 2-s bins by the total EEG power averaged across the recording session.

### Optogenetic manipulation

All optogenetic stimulation were conducted unilaterally. Mice were habituated in the behavioral chamber for at least 4 hours before the experiment. Light pulses (20 Hz, 10 ms) with different durations (30 s, 1 min, 2 min) from a 473 nm laser diode (Shanghai laser & Optics Century Co., Ltd.) were controlled by a microcontroller board (Arduino Mega 2560, Arduino). Laser power is set to 4-6 mW for somatic stimulation in the VLM and 10-15 mW for terminal stimulation in the VLPO. We used two methods to trigger laser stimulation: 1) wake-trigger stimulation: An IR-camera (30 fps) was placed on the ceiling to videotape animal behavior. A custom Matlab program was used to real-time process video frames to detect animal’s location by subtracting each frame from the pre-acquired background image (without the mouse). Laser stimulation was automatically triggered when animal movement continued for a period (5 min). A minimal interval of 30 min was set between trials. In no-light control experiments, the same trigger method was applied except the laser power was off. 2) fixed interval stimulation: inter-stimulation interval for optogenetic stimulation is fixed to 45 min.

### Chemogenetic manipulation

After habituation for 12 h in the testing chamber, C57BL/6J mice expressing hM4Di or hM3Dq in the VLM were injected with saline (day 1) and CNO (day 2, 1 mg/kg body weight) intraperitoneally (i.p.) at the same time of the days. Injections were performed in light cycles (10:00 AM) for chemogenetic inhibition and in dark cycles (9:30 - 10:00 PM) for chemogenetic activation. In control experiments, wild type mice without viral injection were treated with CNO and saline either in light cycles or dark cycles. Sleep recording started at least 1 h before saline injection and lasted 24 h after CNO injection. EEG and EMG in the time window (0.5 – 6.5 h after CNO or saline injection) were used for data analysis.

### Whole brain clearing and imaging

To identify neurons projecting to the POA, we unilaterally injected ∼200 nl AAVrg-hSyn.Cre.WPRE.hGH in the VLPO of Ai9 reporter mice. Six weeks after viral injection, mice were perfused with phospho-buffered saline (PBS) containing 10 U/ml heparin, followed by 4% paraformaldehyde. Brains were then harvested and post-fixed in 4% paraformaldehyde for 3 h at room temperature. The whole brains were cleared by the CUBIC method as previously described(Susaki et al., 2014; Wang et al., 2018). Briefly, mouse brains were washed 3 times in PBS before immersion in CUBIC reagent-1 (diluted 1:2 in water) overnight, incubated in reagent-1 for 7-10 days. Then, brains were washed with PBS, degassed in PBS overnight, and immersed in reagent-2 (diluted 1:2 in PBS) for 6-24 h before incubated in reagent-2 containing TO-PRO-3 (1:5,000, Thermo Fisher Scientific) for additional 7-10 days. Reagent 1 contained 25 wt% urea (Sigma-Aldrich), 25 wt% N,N,N′,N′-tetrakis (2-hydroxypropyl) ethylenediamine (Sigma-Aldrich) and 15 wt% Triton X-100 (Nacalai Tesque). Reagent 2 contained 50 wt% sucrose (Sigma-Aldrich), 25 wt% urea, 10 wt% triethanolamine (Sigma-Aldrich) and 0.1% (v/v) Triton X-100. All clearing procedures were performed at room temperature with gentle shake to prevent sample deformation caused by temperature fluctuation and fluorescence loss. Reagent 1 and Reagent 2 were refreshed every 3 days. Samples were imaged in an oil mix (mineral oil and silicone oil 1:1) horizontally from ventral to dorsal by Light-sheet fluorescence microscopy (UltraMicroscope, LaVision BioTec) as previously described(Wang et al., 2018). The images were acquired with a z-step size of 5 μm. Exposure time was 50 ms per channel per z step. Data was processed in ImageJ (Fiji distribution). The gamma value of the images was set to 0.5 for display purposes. The whole-brain data was registered to a reference atlas (Allen Brain Institute, 25-um resolution volumetric data with annotation map, http://www.brain-map.org) using elastix (version 5.0) (Klein et al., 2010; Shamonin et al., 2013). The voxel size of both sample data and reference template were scaled to 6.5 μm. The 3D reconstruction, cell tracing, and structure labelling were performed in Imaris (version 9.6, Bitplane).

## Statistics

No statistical methods were used to predetermine sample size, and investigators were not blinded to group allocation. No method of randomization was used to determine how animals were allocated to experimental groups. Mice in which post hoc histological examination showed viral targeting or fiber implantation was in the wrong location were excluded from analysis. Paired t test and Mann-Whitney *U*-test were used and are indicated in the respective figure legends. All analyses were performed in MATLAB. Data are presented as mean ± s.e.m.

## Code availability

Custom scripts for EEG/EMG and behavioral analysis are available from the corresponding author upon reasonable request.

## Data availability

All data supporting the findings of this study are available from the corresponding author upon reasonable request.

## Notes

### Competing Interest Statement

The authors have declared no competing interest.

